# Functional organization of visual responses in the octopus optic lobe

**DOI:** 10.1101/2023.02.16.528734

**Authors:** Judit R. Pungor, V. Angelique Allen, Jeremea O. Songco-Casey, Cristopher M. Niell

## Abstract

Cephalopods are highly visual animals with camera-type eyes, large brains, and a rich repertoire of visually guided behaviors. However, the cephalopod brain evolved independently from that of other highly visual species, such as vertebrates, and therefore the neural circuits that process sensory information are profoundly different. It is largely unknown how their powerful but unique visual system functions, since there have been no direct neural measurements of visual responses in the cephalopod brain. In this study, we used two-photon calcium imaging to record visually evoked responses in the primary visual processing center of the octopus central brain, the optic lobe, to determine how basic features of the visual scene are represented and organized. We found spatially localized receptive fields for light (ON) and dark (OFF) stimuli, which were retinotopically organized across the optic lobe, demonstrating a hallmark of visual system organization shared across many species. Examination of these responses revealed transformations of the visual representation across the layers of the optic lobe, including the emergence of the OFF pathway and increased size selectivity. We also identified asymmetries in the spatial processing of ON and OFF stimuli, which suggest unique circuit mechanisms for form processing that may have evolved to suit the specific demands of processing an underwater visual scene. This study provides insight into the neural processing and functional organization of the octopus visual system, highlighting both shared and unique aspects, and lays a foundation for future studies of the neural circuits that mediate visual processing and behavior in cephalopods.

**Highlights:**

- The functional organization and visual response properties of the cephalopod visual system are largely unknown
- Using calcium imaging, we performed mapping of visual responses in the octopus optic lobe
- Visual responses demonstrate localized ON and OFF receptive fields with retinotopic organization
- ON/OFF pathways and size selectivity emerge across layers of the optic lobe and have distinct properties relative to other species

## Introduction

Cephalopods evolved large and complex brains independently from the rest of the animal kingdom. Like vertebrates, cephalopods also evolved camera-type eyes that focus a high resolution image onto a retina (Packard 1972). Together, their large brain and camera-type eyes implement a sophisticated visual system, which mediates a wide range of advanced visually-based behaviors (Hanlon and Messenger 2018), including prey capture and predator avoidance (Schnell et al. 2016; Bidel, Bennett, and Wardill 2022), identifying mates (Shashar, Rutledge, and Cronin 1996; Hanlon et al. 2005), spatial navigation (Karson, Jean, and Hanlon 2003; Alves, Boal, and Dickel 2008), and a remarkable ability to rapidly camouflage in their surroundings (Chiao et al. 2013; Nagar et al. 2021; Reiter and Laurent 2020). However, because the cephalopod brain evolved independently from that of other highly visual species, the neural organization of their visual system is dramatically different. To date, the cephalopod visual system has been largely unexplored at the level of neural function. In contrast to decades of work on visual neuroscience in traditional model organisms, there has not even been a basic characterization of visual processing in the cephalopod central brain, limiting our understanding of the neural computations that mediate vision in their unique and complex brains.

Anatomical studies have delineated the morphology and structural connectivity of neurons in the cephalopod retina and optic lobes (Young 1971, 1960; Ramón y Cajal 1930) and identified connections from the optic lobe to other areas of the brain (Saidel 1982; Chung, Kurniawan, and Marshall 2020; Williamson and Chrachri 2004). Unlike those of vertebrates, cephalopod retinas are relatively simple, containing only photoreceptors and a small population of presumptive horizontal cells that connect them. The photoreceptors themselves send axons out of the retina into the optic lobes (Figure 1A, B), which comprise roughly two thirds of the centralized nervous system and are where most of the visual processing in the cephalopod brain is thought to occur (Wells 1962). The outer optic lobe is a layered structure (Figure 1B, C), with two cell body layers, termed the outer granular (OGL) and inner granular layer (IGL), surrounding a layer of processes, the plexiform layer (Plex), where photoreceptor axons terminate. The central portion of the optic lobe, the medulla (Med), consists of clusters of cell bodies arranged in a tree-like structure surrounded by neuropil (Liu et al. 2017). Neurons of the medulla have processes within superficial layers of the optic lobe as well as locally within the medulla, and many project axons out from the optic lobe to downstream visual areas. Recent transcriptomic studies have further revealed a rich diversity of cell types within the optic lobe, as well as extensive sub-laminar organization (Songco-Casey et al. 2022; Styfhals et al. 2022; Gavriouchkina et al., 2022; Duruz et al. 2023).

**Figure 1.**
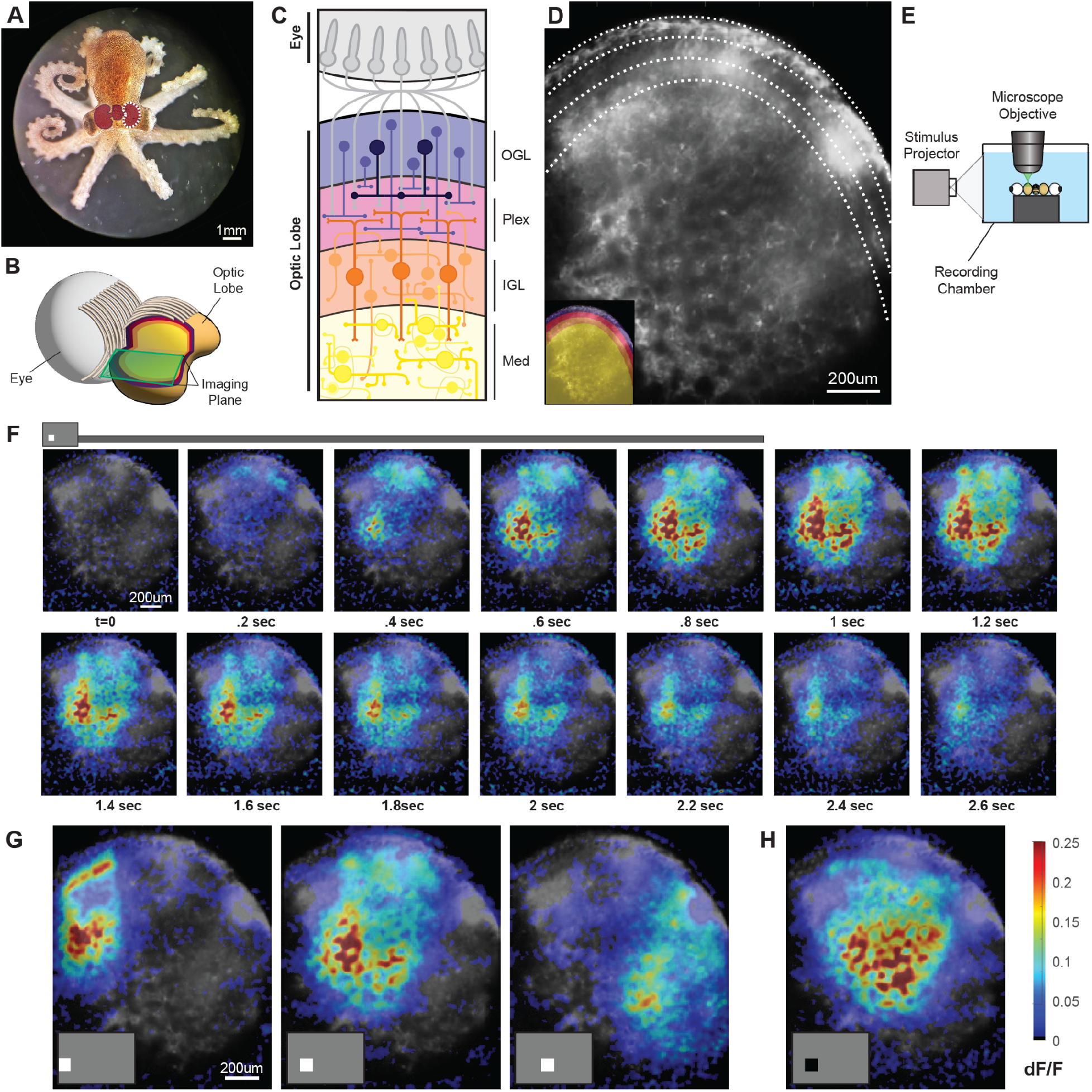
Experimental paradigm for calcium imaging of visual responses in the optic lobe. A) Image of a juvenile *Octopus bimaculoide*s. Between their eyes is the central brain complex, including two optic lobes, one behind each eye. An outline of the central brain is shown in burgundy, and the position of the right optic lobe is indicated by the white outline. B) Illustration of octopus visual system anatomy. Bundles of photoreceptor projections exit the back of the eye (left), decussate vertically, and then enter the optic lobe (right) in a retinotopic manner. In the cutaway, the layered structure of the optic lobe can be seen, as it is in our imaging planes. C) Simplified illustration of the anatomy of the layers of the optic lobe. Photoreceptors in the eye send projections through a cell body layer on the surface of the optic lobe (outer granular layer, OGL, purple) to a layer of neuropil below it (plexiform layer, Plex, red). Here most photoreceptors synapse onto projections from cells in the layer below (inner granular layer, IGL, orange). These in turn send projections to the interior of the optic lobe (medulla, Med, yellow). Color code for layers also applies to Figure 1B,D. D) Mean fluorescence image of Cal-520 calcium indicator loading in the optic lobe, demonstrating successful labeling across multiple layers, delineated by dotted lines. Inset shows layers in color overlay. E) Schematic of the experimental set-up. A projector is used to present visual stimuli on the side of the recording chamber, with the preparation underneath the objective of a two-photon microscope. F) Mean timecourse of fluorescence response to a flashed ON spot at one location in the visual field (averaged over five stimulus repetitions), showing spatial organization and temporal dynamics. Stimulus onset and offset are indicated in the gray bar below the frames, and individual frames are shown at 0.2sec intervals. G) Mean fluorescence response across the optic lobe to ON stimuli at three different horizontal locations, averaged across the stimulus duration for five repetitions. H) Mean fluorescence response to an OFF stimulus at the same location as G (middle), averaged across the stimulus duration for five repetitions.

Early studies of photoreceptors in the cephalopod eye provided an initial description of visual processing at the input stage (Hamasaki 1968; Lange and Hartline 1974). Like most other invertebrates (Land and Fernald 1992), cephalopods have rhabdomeric photoreceptors that depolarize in response to increases in light (ON responses) (Moccia, Cristo, and Di Cosmo 2009), in contrast to vertebrate photoreceptors that depolarize in response to decrements in light (OFF responses). Nearly all species of cephalopods, including octopuses, only express one type of opsin in their photoreceptors and are therefore thought to be colorblind (Hamasaki 1968). Electrophysiological recordings of photoreceptor visual responses have demonstrated ON-center receptive fields and indications of lateral inhibition (Tasaki, Oikawa, and Norton 1963; Norton et al. 1965; Hartline and Lange 1974; Saidel et al. 2005). In contrast, little is known regarding neural responses beyond the photoreceptors. A small fraction of optic lobe neurons project back to the retina, and recordings from the optic nerves have shown these to have slow responses and large receptive fields (Hartline and Lange 1974; Patterson and Silver 1983). Bulk field potential recordings in the optic lobe of freely moving octopuses demonstrated oscillatory responses to brief light flashes, which displayed intriguing state-dependence (Boycott et al. 1965).

However, no studies have addressed the neural representation of visual stimuli within the optic lobe, or how this is organized topographically and transformed across the optic lobe layers. This information is an essential foundation for understanding how neural pathways in the cephalopod brain process visual scenes. Here we developed techniques for two-photon calcium imaging of visually evoked responses in *Octopus bimaculoides* (Pickford and McConnaughey 1949), a promising model species for studying cephalopod vision (Albertin and Simakov 2020). We used this calcium imaging approach to measure how spatial and luminance information are represented in large-scale neural responses, and to determine how these responses are organized within the optic lobe.

## Results

### Calcium imaging of stimulus-specific visual responses in the optic lobe

Historically, electrophysiological recordings in the cephalopod brain have been technically challenging, and methods to express genetically encoded calcium indicators are not yet available in cephalopods. Here instead we employed a calcium imaging approach based on injection of a synthetic calcium indicator, Cal-520 AM-ester, to measure large-scale neural activity in the octopus optic lobe. Our general approach for calcium imaging and visual stimulation is adapted from techniques previously used to measure visual responses in the zebrafish optic tectum (Niell and Smith 2005), and loading methods established for AM-ester calcium indicators (Garaschuk et al. 2006). Briefly, we injected Cal-520 into one optic lobe of an ex vivo preparation comprised of the eyes and central brain of an octopus. We imaged neural responses with a two-photon microscope, which provided optical access for recording across the optic lobe at depths of 100-200um. The small sizes of the juvenile octopuses allowed us to image a large cross-section of their optic lobes spanning multiple layers in a single field of view. The optic lobe is a three-dimensional structure similar to a flattened sphere, so a given optical section from two-photon imaging at this depth provides a view across both its tangential and laminar organization (Figure 1B, C). Figure 1D shows loading of the fluorescent indicator across an optic lobe, with its different layers readily discernible (Figure 1D). Controlled visual stimuli were displayed via a LCD projector onto a white diffusion filter mounted on the side of the chamber containing the preparation (Figure 1E). An adjustable platform allowed us to center one eye’s field of view on the screen, and recording was performed on the corresponding optic lobe. This approach allowed us to present high-resolution stimuli across the visual field of one eye while simultaneously recording the responses across the optic lobe.

To obtain visually evoked responses, we initially used a stimulus consisting of full contrast ON and OFF rectangular spots (24×18 deg) on a 6×4 grid spanning the projection screen, presented in a random order for a one second duration. This stimulus elicited fluorescence responses in the optic lobe dependent on the location of the spot in the visual field (Figure 1F-H, and Supplemental Video 1). Figure 1F shows the mean response, measured as the change in fluorescence divided by mean fluorescence (dF/F) at each pixel across the optic lobe, over five repeated presentations of an ON spot at one location. The evoked activity, locked to stimulus onset, persisted throughout the one second stimulus period and was followed by a decay, consistent with calcium indicator dynamics. This activation map also suggests a temporal sequence of activity, with fluorescence signal first increasing rapidly in the superficial optic lobe, followed by more gradual and sustained response in the medulla. Figure 1G shows the mean response across the optic lobe during the stimulus presentation for ON spots in three neighboring locations. We found activation of distinct regions within the optic lobe to each location, indicating specificity for the location of the stimulus in visual space in a retinotopic manner. Finally, Figure 1H shows the response to an OFF spot at the same recording location as Figure 1G (middle), demonstrating a response in approximately the same region, but deeper in the laminar structure of the optic lobe, in the IGL and medulla.

These results demonstrate that our calcium imaging approach allowed us to measure stimulus-specific visual responses, and provide initial support for both retinotopic and laminar organization of responses. To probe the specificity and spatial organization of visual responses more systematically, we next performed mapping of spatial receptive fields using a sparse noise stimulus.

### Spatially localized ON and OFF receptive fields

We used a sparse noise stimulus adapted from (Piscopo et al. 2013) to calculate ON and OFF receptive fields. The stimulus consisted of frames of ON and OFF circular spots of three different sizes (radius = 3, 6, 12 deg) in a randomly distributed pattern, along with randomly interleaved ON or OFF full-field frames (Figure 2A). Each frame was presented for 1sec over a total recording time of 10mins. This sparse noise stimulus elicited robust and spatially localized fluorescence responses across the optic lobe, as demonstrated in Figure 2B and Supplemental Video 2.

**Figure 2.**
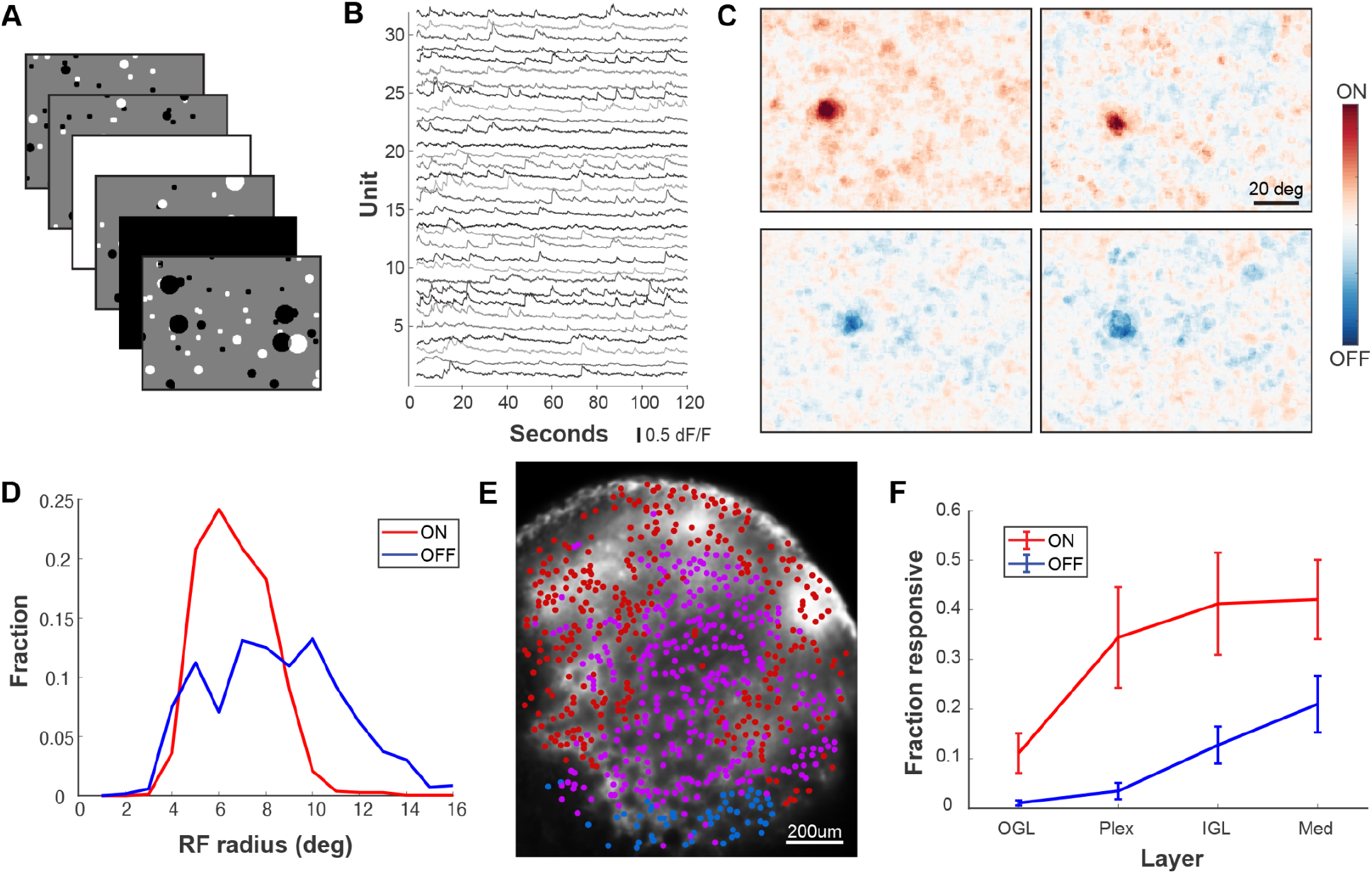
ON and OFF receptive fields mapped with a sparse noise stimulus. A) Example frames from the sparse noise stimulus used to map receptive fields. Frames were presented consecutively in a randomized order for a 1sec duration each. B) Traces of fluorescence activity at 32 locations across the optic lobe recorded in response to the sparse noise stimulus. C) RFs from four example units, two each for ON (top) and OFF (bottom) components of the stimulus. D) Histogram of RF sizes for ON and OFF stimuli (N=6 experiments). E) Location of units with RFs for ON (red), OFF (blue), or both (magenta) in one session across the optic lobe. F) Fraction of units overall with significant RFs for ON and OFF across the layers of the optic lobe (N=6 experiments).

For analysis, we selected individual regions of interest (ROIs), 20×20μm, centered on peaks in the mean fluorescence image that were above a baseline fluorescence threshold, to exclude regions that were not loaded with the calcium indicator. This identified ∼500-1000 ROIs spaced across each of the multiple layers of the optic lobe captured within each imaging field (e.g Figure 2E). We selected this approach, rather than extracting responses specifically from cell bodies as typically performed for calcium imaging in vertebrates, both due to the challenge in localizing signals to individual cells in tightly packed cell body layers and the fact that, in invertebrates, much of the neural signal is localized to processes within the neuropil. We refer to each ROI as a unit, denoting a specific location within the optic lobe, rather than a single neuron. This analysis allowed us to map how visual information is represented at locations across the optic lobe. As noted in the Discussion, single-cell or cell-type specific recordings will likely be needed to directly probe individual cell tuning properties.

We computed receptive fields (RFs) for each unit using reverse correlation based on the evoked dF/F fluorescence signal for each frame of the sparse noise stimulus, excluding the full-field flashes (see Methods). We performed this separately for the ON and OFF components of the stimulus to avoid cancelation of positive and negative stimulus contrast for units that responded to both polarities. This revealed spatially localized RFs for both ON and OFF stimuli, as shown by examples in Figure 2C. We fit RFs to a Gaussian model to determine their size and location within visual space. Across experiments, 59 +/- 26% of all units had a RF significantly above background as determined by their z-scored response. The RF radius, based on sigma of the Gaussian fit, was 5.7 +/- 0.6 deg for ON, and 7.4 + /- 0.6 deg for OFF (p=0.31 for ON vs OFF, N=6 experiments) (Figure 2D). Note that this is likely an overestimate of the RF size of individual neurons, since the response of each unit within the lobe represents the summed response of a number of individual neurons.

We next examined the distribution of ON and OFF responses across the optic lobe to determine where the pathway for processing each arises. Figure 2E shows all units in an example recording labeled based on whether they had a RF for only the ON (red) component of the stimulus, only the OFF (blue) component, or for both (magenta). This demonstrates that while ON responses are distributed throughout the lobe, OFF responses are largely restricted to the deeper layers of the IGL and medulla. To quantify this, we calculated the fraction of ON and OFF RFs in each layer across recordings (Figure 2F), confirming that OFF responses primarily emerge in the IGL and are strongest in the medulla. The sequential emergence of OFF responses relative to ON is consistent with the fact that photoreceptor axons in cephalopods, which mainly terminate in the superficial layers of the optic lobe (Plex), respond to increments of light, and demonstrates that the OFF processing pathway in octopuses likely originates in neurons further along the visual processing pathway.

### Retinotopic organization of the optic lobe

To determine if there was a retinotopic organization of visually evoked responses in the octopus central brain, we labeled each unit according to the location of its RF, based on the center of the Gaussian fit described above. As shown in Figure 3A, we found clear retinotopic progression for ON and OFF responses, along both the elevation and azimuth axes of the visual field, resulting in a map of visual space across the optic lobe. This is further demonstrated in Figure 3B, which shows the high degree of correlation between RF location in visual space with the unit’s physical location across the optic lobe. The retinotopic maps of ON and OFF RFs were also aligned in regions of the lobe that were responsive to both (Figure 3B).

**Figure 3.**
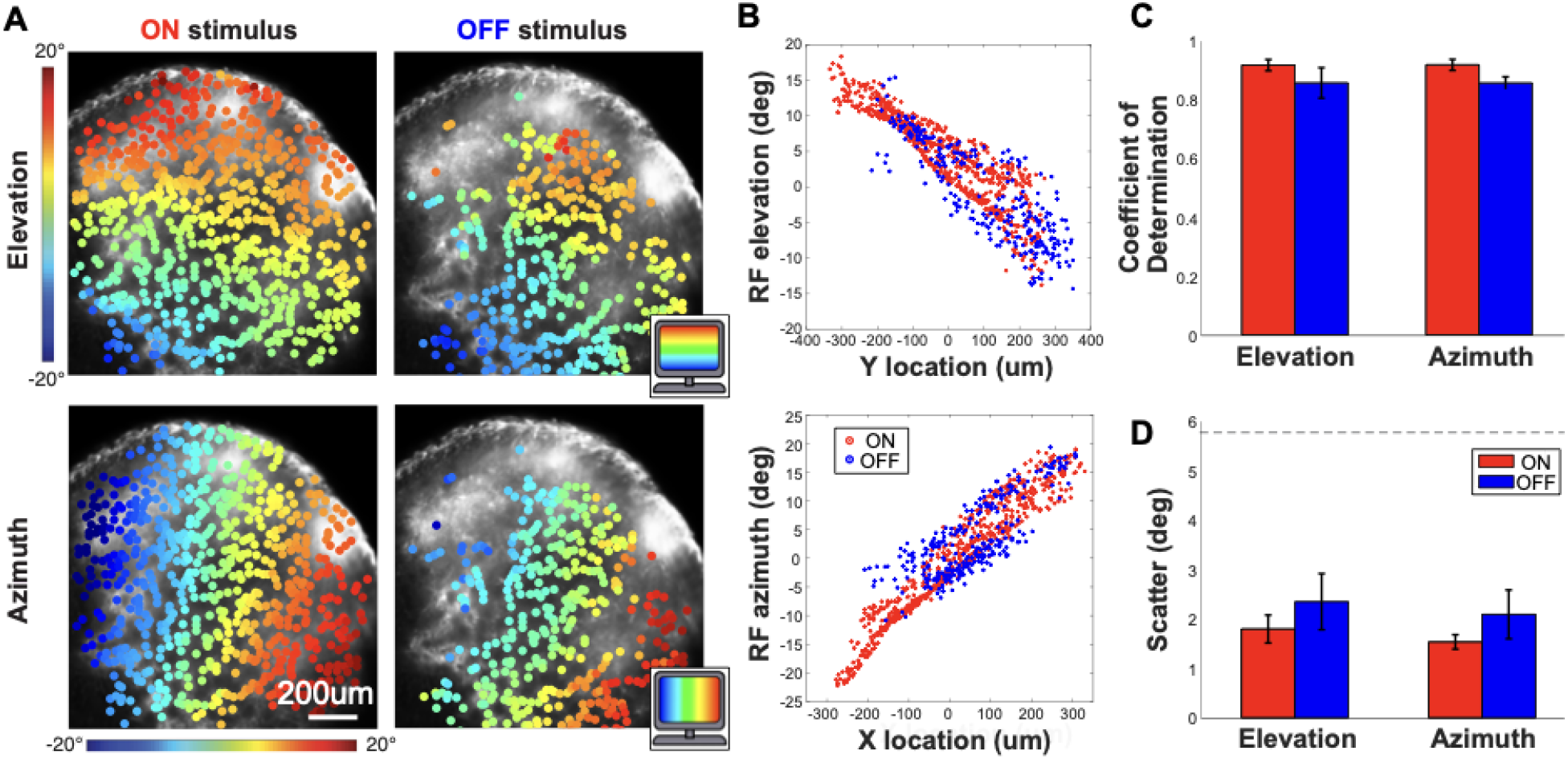
Retinotopic organization of visual responses in the octopus optic lobe. A) Example mapping of RFs in the optic lobe of responses to both ON (left) and OFF (right) stimuli. Areas are colored by the position of their RFs along the elevation (top) and azimuth (bottom) as shown by the color scale bars (degrees). B) Scatter plot of RF location for elevation (top) and azimuth (bottom) versus unit location within the optic lobe, for both ON and OFF responses. Adjacent groups of cells responded to adjacent areas of the visual field. C) Mean coefficient of determination for elevation and azimuth maps across all recordings (N=6 experiments). D) Mean scatter in RF location for elevation and azimuth, across all recordings (N=6 experiments). Dashed line shows chance level based on a shuffle control.

We next quantified the retinotopic organization in each experiment by performing a linear regression between the RF elevation/azimuth in the visual space of all responsive units and their x/y location within the optic lobe. Note that we used both x and y location of the units to predict each RF parameter, since the visual axes were not always aligned to the x and y axes of the imaging plane depending on the orientation of the preparation. This fit resulted in a mean coefficient of determination greater than 0.8 for both ON and OFF maps across experiments (Figure 3C), confirming robust retinotopy. We also computed the scatter of RF locations (i.e. how much RF locations deviate from a linear retinotopic progression), based on the residuals from the fit, which demonstrated that individual unit’s RFs have scatter of less than 2 degrees (Figure 3D). Finally, the slope of the RF fit determines the magnification factor of the map (i.e. how much the RF location changes for a given distance in the brain), with a mean progression of 21.9 +/- 1.4 um/deg in elevation and 25.0 +/- 3.0 um/deg in azimuth. Together, these data provide the first functional demonstration of a retinotopic organization of visual information within the cephalopod brain.

### Size selectivity in ON and OFF pathways across layers of the optic lobe

To further examine visual response properties and their organization within the octopus optic lobe, we next calculated size tuning of units, based on their evoked responses to spots of different sizes in the sparse noise stimulus. For each unit with a significant RF, we determined the center of its ON or OFF RF from the Gaussian fit, and computed the mean dF/F response timecourse when spots of different sizes appeared at this location during the stimulus. We limited this analysis to units with significant RFs, based on z-score as described above, because it is only meaningful for units that have a defined RF location.

Figure 4A shows the mean timecourse response to each size stimulus, including full-field flash, for all units across experiments, according to their layer within the optic lobe. In order to accurately represent the relative magnitude of responses across layers, given the differential distribution of ON and OFF units (Figure 2E), we weighted these mean traces by the respective fraction of responsive units within each layer.

**Figure 4.**
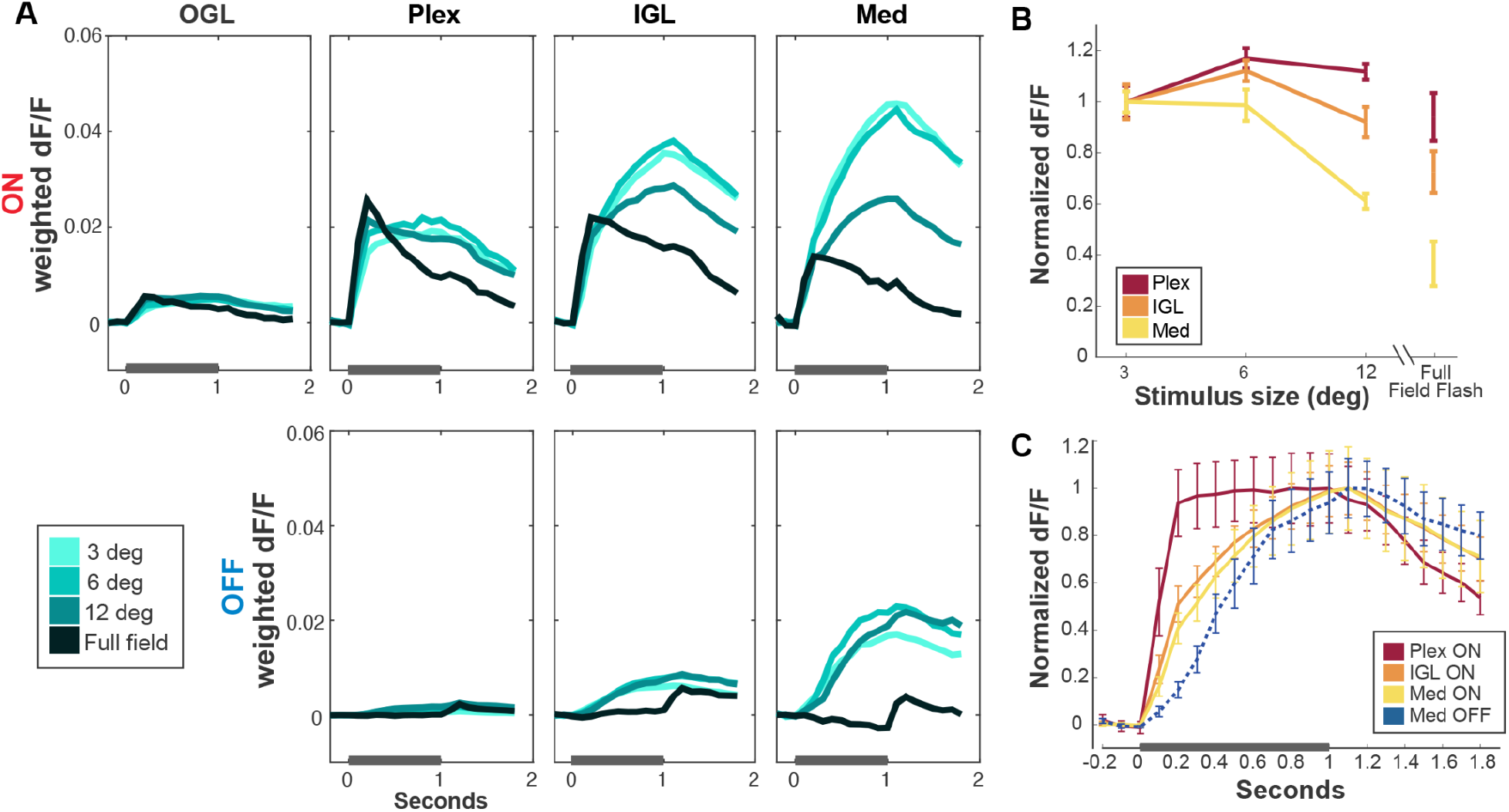
Size selectivity and temporal dynamics across the layers of the optic lobe. A) Mean timecourse of ON (top panels) and OFF (lower panels) responses for each stimulus size, separated by layers of the optic lobe. Response for each layer and luminance are weighted by fraction of units responsive. OGL did not show a significant response to OFF, and therefore was omitted from this figure. Stimulus onset is at t=0 and each frame was presented for 1sec, as shown by gray bars on the x axis (N=6 experiments). B) Mean size tuning curves for ON responses in each layer, normalized to the response to the smallest stimulus (N=6 experiments). C) Mean timecourse of unit responses, averaged across the three sizes of stimulus spots and normalized to the maximum response, for ON (Plex, IGL, Med) and OFF (Med) (N=6 experiments).

For ON stimuli (Figure 4A, top), there was a strong and rapid response in the Plex, which was approximately equal in amplitude for all stimulus sizes, as well as to the full-field flashes. However, as responses progressed deeper into the IGL and medulla, the sustained response increased for small stimuli but decreased for larger spots and full-field flashes, indicating size selectivity. Interestingly, the initial onset responses were similar across sizes, with the responses to different sizes diverging only after ∼200msec. Figure 4B shows the size tuning curve of ON responses for each layer, based on the mean dF/F across stimulus duration, normalized to the response to the smallest stimulus size. These show a decrease in the relative response to larger stimuli in the IGL and medulla. Together, the responses to ON stimuli therefore demonstrate an emergence of size selectivity, both over time and across layers.

In contrast, responses to OFF stimuli (Figure 4A, bottom) only appear in deeper layers of the optic lobe (IGL and Med). There was no size suppression across different sized spots in OFF response, in contrast to what was seen in the ON. Rather, responses to OFF spots of all sizes were roughly equal, leading to a relative bias toward large stimuli in OFF compared to ON. Strikingly, there was no response at all to the full-field OFF stimulus, despite responses to the range of OFF spot sizes and to full-field ON. These differences in spatial integration for ON and OFF suggest different processing pathways exist for these two luminance modalities, even in these early visual processing stages of the optic lobe.

Finally, we examined the mean timecourse of responses to ON and OFF spots across the layers of the optic lobe (Figure 4C), revealing distinct temporal dynamics. ON responses emerged first in the plexiform layer, and then spread into the deeper regions of the IGL and Med. On the other hand, OFF responses were first seen predominantly in the medulla (Figure 4A, bottom panels), and showed a slower rise time in response (Figure 4C, blue trace), consistent with a later emergence in the visual processing circuit.

## Discussion

Octopuses represent an intriguing independent evolution of a complex nervous system. However, relatively little is known about how their brain functions at the neural level. Combining large scale two-photon calcium imaging with the presentation of controlled visual stimuli, we were able to overcome technical challenges that previously hindered recordings of neural activity in cephalopods. The establishment of such recording techniques, and future improvements, will be essential for elucidating the computations performed in the visual system, as well as other aspects of sensory processing, cognition, and behavior in cephalopods.

Using the calcium imaging approach we measured response properties of populations of neurons within the octopus optic lobe, and began to identify what fundamental features of the visual world they encode, and how these emerge in the early stages of visual processing. We found similarities in visual processing between octopus and other species, such as a retinotopic organization of responses, highlighting potential fundamental principles for the organization of visual systems across the animal kingdom. We also identified differences, such as the organization of ON/OFF pathways and size selectivity, that may have arisen due to these animals’ unique environmental constraints (Chung and Marshall 2017) or distinct evolutionary trajectories (Grasso and Basil 2009). These findings provide the first demonstration of visual processing dynamics across the layers of the octopus optic lobe and provide a foundation for studying the processing of more complex visual features.

### Spatial organization of response properties in the optic lobe

Although there have been previous studies of the anatomical organization of the octopus visual system, little is known about its functional organization. Based on the orderly fashion in which the optic nerves from the eye were found to enter the optic lobe (Young 1971; Saidel 1979; Chung, Kurniawan, and Marshall 2022), it was predicted that visual information would be retinotopically organized within the lobe, as it is in many, though not all (Hoseini et al. 2018), visual systems across the animal kingdom. However, studies in the motor system of cephalopods have demonstrated a surprising lack of somatotopy in their central brain, leading to the suggestion that they may have evolved alternative, non-topographic architectures for representing spatial information (Zullo et al. 2009). In this study, we found that neural coding in the visual system of the octopus is indeed organized retinotopically, with aligned maps for responses to ON and OFF stimuli. This demonstrates that the lack of topographic organization observed for somatotopy is not a general feature of cephalopod brain organization.

Previous anatomical studies had suggested potential neural circuits across the layers of the octopus optic lobe that could implement sequential processing of visual input (Ramón y Cajal 1930; Young 1971), as in the vertebrate retina or fly visual system (Sanes and Zipursky 2010). Our findings support these predictions, demonstrating that the temporal dynamics of visual responses in octopuses do in fact proceed sequentially across the laminar organization of their brain. This is accompanied by a transformation of the visual input including the emergence of the OFF pathway, as well as an increase in size selectivity in the ON pathway. These findings of differential response dynamics across distinct layers provide an initial framework for understanding the functional computations performed by the circuitry of the cephalopod visual system.

### Comparative aspects of ON/OFF pathways and spatial processing

A key computation for any visual system is the ability to respond to both light and dark stimuli within a scene. Given that photoreceptors depolarize to only ON (invertebrates) or OFF (vertebrates) stimuli, there is a necessary computation to invert the polarity of the photoreceptor signal within the subsequent visual circuitry to do so. For vertebrates, it is known that this inversion occurs at the photoreceptor to bipolar cell synapse, where ON and OFF bipolar subpopulations segregate the response to light and dark stimuli coming from the retina in the initial stages of visual processing. Since both pathways receive input directly from the same photoreceptors, the information entering ON and OFF pathways is similar but of an opposite sign. In Drosophila, segregated ON and OFF responses emerge one synapse further from the photoreceptors, between the lamina and medulla, with lamina L1 neurons inhibiting their postsynaptic partners while lamina L2 neurons excite their postsynaptic partners (Behnia et al. 2014).

Here we found that ON responses dominate in the primary input layer of the octopus optic lobe, the plexiform layer, corresponding to the fact that cephalopod photoreceptors depolarize to light increments. This is striking, as the plexiform layer also contains processes from many cell types in addition to photoreceptor terminals. OFF responses only emerge initially in the IGL and are greatly increased in the medulla, suggesting a potential site for the sign inversion needed to translate the response from photoreceptor inputs that depolarize in response to light. We also found that OFF responses have a strikingly different profile to those of ON responses, in that they completely lack a response to full-field OFF stimuli, despite prominent responses to full-field ON stimuli across layers. This suggests that the OFF pathway may emerge through a different mechanism than direct inversion of the photoreceptor input, which would yield responses to a full-field OFF stimulus. One possibility is that the OFF pathway receives input from a subset of ON neurons that have completely suppressed the response to a full-field stimulus. A more intriguing possibility is that the mechanisms that generate OFF responses may rely directly on boundaries between light and dark regions, which would explain why OFF responses are driven by localized dark stimuli (i.e spots) that contain such edges, but not full-field stimuli, which do not.

In addition, we also found differences in size selectivity for spots in the ON and OFF pathways (Figure 4A). While responses in the ON pathway decreased for larger spots, the responses to spots in the OFF pathway were roughly equal across the sizes of stimuli we measured. This implies a net bias toward the enhancement of responses to smaller stimuli in the ON pathway. Asymmetries in ON/OFF visual processing pathways have been found in other species across the animal kingdom, and are thought to enhance ethologically relevant visual features to meet each animals’ specific visual demands. In the early stages of processing in the Drosophila visual system, there are differences in the temporal response to ON and OFF moving edges, which are thought to be driven by the statistics of natural scenes (Leonhardt et al. 2016; Clark et al. 2014). In the cat visual cortex, ON and OFF responses seem to have evolved to favor global-slow and local-fast stimuli respectively, which is matched to their visual environment, and has been proposed to lead to specialized roles in image stabilization versus high acuity vision (Mazade et al. 2019).

It is interesting that the asymmetry in ON/OFF processing we found in the octopus is the opposite of what has been found in vertebrates (Mazade et al. 2019). The enhancement of responses to smaller stimuli in the ON pathway may be beneficial when processing visual scenes underwater, where light intensity is greatly attenuated by both absorption and scatter (Cronin et al. 2014). As a result, nearby objects will tend to appear bright against a large, dark background. An OFF pathway biased towards larger stimuli might also aid in the detection of large, looming objects, which often represent predators. It will be interesting to see if such ON/OFF processing differences exist more broadly across cephalopods that occupy other ecological niches, particularly as these vary greatly in luminance levels and visual scene statistics (Chung and Marshall 2016).

### Implications for future studies

Our findings provide initial insight into how luminance and size are processed at the level of layers within the octopus optic lobe. However, both anatomical and transcriptomic studies have revealed a high degree of cell type diversity within these layers, so the bulk response properties we examined here undoubtedly mask a high degree of underlying functional diversity. Identifying the detailed response properties within the parallel pathways of diverse cell types in this system will likely require novel methods to record from genetically identified cells, not yet available in cephalopods to date. While it may be possible to achieve this using post-hoc identification of cell type identity (Kerlin et al. 2010), a more promising approach would be the use of cell-type specific expression of genetically encoded indicators. Such an approach will also help address the challenge in associating activity in neural processes, which often dominate in invertebrate neurons, with individual neurons or populations of neurons. This has been used to dissect visual selectivity and pathways in the Drosophila visual system (e.g. Strother, Nern, and Reiser 2014), even where signals are intermingled in the neuropil.

More broadly, future studies based on these findings and methodology could explore a broad range of feature selectivity in octopuses, as has been studied in other species, such as motion processing, orientation selectivity, object recognition, and lateralization of visual responses (Byrne, Kuba, and Meisel 2004; Frasnelli et al. 2019). Additionally, this approach can be used to study aspects of visual responses that may be specific to cephalopods, such as the ability to detect stimuli based on the polarization angle of light (Shashar 2014), or to extract information from the visual scene for camouflage (Reiter and Laurent 2020). It also remains unknown how the neural circuits of the cephalopod visual system are assembled across development to establish these response properties (Liu et al. 2017). Further measurement of visual response properties, alongside methods for circuit tracing and manipulations of neural activity, may reveal how the cephalopod brain performs the computations that enable the remarkable visual capabilities of these enigmatic creatures.

## Acknowledgements

We thank members of the Niell lab past and present for helpful discussions and comments on the manuscript, and members of the Hochner lab (Hebrew University of Jerusalem) for advice on experimental methods for octopus. We would also like to thank Rhanor Gillette, Spencer Smith, and Michael Wehr for feedback on the manuscript. This work was supported by National Institutes of Health R01NS118466-01, Office of Naval Research N00014-21-1-2426, and Human Frontiers Science Program RPG0042/2019.

## Author Contributions

J.R.P. and C.M.N. conceived the project and designed experiments. J.R.P. led the project and performed experiments. V.A.A. and J.O.S.-C. both optimized the experimental protocol and performed experiments, contributing equally. C.M.N. and J.R.P performed data analysis. All authors contributed to the writing of the manuscript.

## Supplementary Information

Supplementary Video 1: Example video of calcium imaging during presentation of 16×24deg ON and OFF spots, corresponding to Figure 1. Time-locked stimulus is shown in upper right, with the dashed rectangle delineating the approximate region of visual space eliciting responses in this field of view. Video is presented at 3X real-time. Scale bar equals 100um.

Supplementary Video 2: Example video of calcium imaging during presentation of the sparse noise stimulus, corresponding to Figure 2. Time-locked stimulus is shown in upper right, and video is presented at 3X real-time. Scale bar equals 100um.

## Methods

### Animal use and husbandry

All studies were conducted with approved protocols from the University of Oregon Animal Care Services, in compliance with the Association for Assessment and Accreditation of Laboratory Animal Care International guidelines. Animal husbandry and protocols were carried out in accordance with published guidelines for the care and welfare of cephalopods in the laboratory (Fiorito et al. 2015, 2014).

*Octopus bimaculoides* were obtained from the Cephalopod Resource Center at the Marine Biological Laboratory (Woods Hole, MA) and from Aquatic Research Consultants (Dr. Charles Winkler, San Pedro, CA). Animals used were 4-8 weeks old and of indeterminate gender. Octopuses were kept in a custom built 250 gallon circulating seawater system, held at 22°C and lit on a 12/12hr day/night light cycle. Each animal was kept in an isolated enclosure within the system, allowing for ample freedom to roam, while keeping them isolated from potential cannibalism from counterparts. Each individual housing was equipped with fixed items that provided shelter for animals (large shells, tubes), and varied other items (smaller shells, Legos, beads), one third of which were rotated weekly to provide environmental enrichment. Animals were fed a mixed diet of frozen shrimp, clams, and fish, offered daily.

### Calcium imaging

Animals were deeply anesthetized in artificial seawater (ASW) (460mM NaCl2, 10mM KCl, 10mM glucose, 10mM HEPES, 55mM MgCl2, 11mM CaCl2, 2mM glutamine, pH 7.4) supplemented to contain 110mM MgCl2 at 13-15 °C. When the animal was no longer responsive to a pinch test of the mantle, it was transferred to an oxygenated dish of a 30:70 mix of isotonic 370mM MgCl2 with ASW that was held between 13-15°C. Animals were then rapidly euthanized via decapitation and removal of the arm crown. A solution to dilate the pupils (10% phenylephrine HCl in ASW) was manually applied to the eyes. Dissection was performed to expose the brain and remove musculature to reduce motion artifacts during recording.

The ex vivo preparation of the central brain and eyes was secured to a coverslip using cyanoacrylate (Vetbond, 3M). A dye solution of 1mM Cal-520 AM (AAT Bioquest), 2.5% Alexa Fluor™ 568 Hydrazide (Thermo Fisher), 8% dimethylsulfoxide, and 2% pluronic acid (AAT Bioquest) in ASW was injected into one of the optic lobes using a glass micropipette needle (Harvard Apparatus Cat. Num. 30-0038) using a pressure injector (ASI, Inc). After injection, the preparation was covered in a thin layer of 4% low melt agarose in ASW (Sigma) to secure the preparation and minimize movement. This paradigm was adapted from previous work in zebrafish (Niell and Smith 2005), see also (Koizumi et al. 2018).

The preparation was kept in a recording chamber filled with ASW and continuously oxygenated via an airstone. The recording chamber consisted of a 7.6cm x 7.6cm x 5cm plastic box (TAP Plastics) where one side was replaced with a white diffusing glass (Edmund Optics Cat. Num. 02-149) to serve as a projection screen for visual stimuli. The coverslip with the mounted preparation attached was secured to a custom-built rotatable platform within the recording chamber to allow for alignment of the preparation to the stimulus screen. The eye ipsilateral to the loaded dye was placed 2cm from the screen for recordings. The chamber temperature was monitored and held between 17-22°C.

Calcium imaging was performed with a two-photon microscope (Neurolabware Inc.), using a 16X Nikon CFI75 LWD objective, via the Scanbox software package for Matlab (MATHWORKS). Data were acquired at a 10Hz framerate, with an 800 × 800um (796 × 796 pixel) field of view.

### Visual stimuli

Custom generated visual stimuli, rendered using the PsychToolbox package for Matlab (Brainard 1997), were displayed with a pico LCD projector (AAXA Technologies) onto the diffusing glass on the side of the recording chamber. To avoid light from the stimulus entering the two-photon detection pathway, the projected light was passed through a 450/50 bandpass filter (Chroma Technology Corporation), avoiding overlap with the emission spectrum of the Cal-520 calcium dye. This also coincides with the absorption spectrum of cephalopod photopigments (Hamasaki 1968). RFs were mapped using a sparse noise stimulus, consisting of white and black spots (radius = 3, 6, 12deg; density = 10%) on a gray (50% luminance) background, along with full-field white or black on 2% of frames. Each stimulus frame was presented for 1sec in a randomized order.

### Data analysis

Data analysis was performed using custom software in MATLAB. We applied a rigid alignment of imaging data using the *sbxalign* function in Scanbox (Neurolabware, Inc.). In order to detect large movements that were not corrected by the alignment algorithm, for each frame we calculated the pixel-wise correlation coefficient to the mean image. Frames with less that 90% correlation were discarded from further analyses. To analyze local responses, we defined “units” as a 20μmx20μm wide square window, centered on local peaks within the mean fluorescence that were above the background fluorescence, to ensure that only areas with sufficient dye loading were analyzed. Units were manually assigned to anatomical layers (OGL, IGL, Plex, and Med) based on location within the mean fluorescence image from the recording session.

To analyze receptive fields (RFs), based on the sparse noise stimulus, we first calculated the evoked response, *r*(*t*), for each frame as the mean dF/F across the one second duration of stimulus presentation, minus the mean dF/F in the preceding 300msec. RFs were calculated by reverse correlation between the each stimulus frame, *s*(*x, y, t*), and the evoked response to that frame.

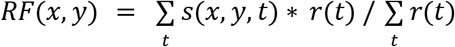

We computed the z-score for each RF based on the maximum absolute value of the RF, divided by the standard deviation across pixels. We used a z-score of 5.5 as the threshold for significant responses.

In order to analyze RF size and location, we fit each RF to a Gaussian function, defined as

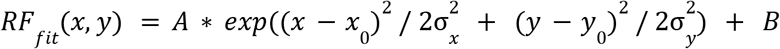

We used *x*_0_, *y*_0_ as the receptive field center, and computed RF radius as (*σ*_*x*_ +*σ*_*y*_)/2. To quantify topographic maps, we performed a linear regression for each recording for responses to both azimuth and elevation, as a function of each unit’s location within the optic lobe from the Gaussian fit, and used the coefficient of determination and standard deviation of residuals (scatter) as metrics of retinotopy.

### Statistics

Statistical tests for comparison of responses across populations within the optic lobe were performed using a t-test. To account for the nested design (many units per recording) of this analysis, all statistical tests were performed based on recordings, rather than total number of units recorded. Summary statistics in text and figures are presented as mean +/- standard error, unless otherwise noted.

